# Neurofilament light as a blood biomarker for neurodegeneration in Down syndrome

**DOI:** 10.1101/281139

**Authors:** Andre Strydom, Amanda Heslegrave, Carla M Startin, Kin Y Mok, John Hardy, Jurgen Groet, Dean Nizetic, Henrik Zetterberg

## Abstract

**INTRODUCTION:** Down syndrome (DS) may be considered a genetic form of Alzheimer’s disease (AD) due to universal development of AD neuropathology, but diagnosis and treatment trials are hampered by a lack of reliable blood biomarkers. A potential biomarker is neurofilament light (NF-L), due to its association with axonal damage in neurodegenerative conditions.

**METHODS:** We measured blood NF-L concentration in 100 adults with DS using Simoa NF-light^®^ assays, and examined relationships with age, and cross-sectional and longitudinal dementia diagnosis.

**RESULTS:** NF-L levels increased with age (Spearman’s rho = 0.789, p<0.001), with a steep increase after age 40, and were predictive of dementia status (p=0.022 adjusting for age, sex, and APOE4) but showed no relationship with longstanding epilepsy or premorbid ability. Baseline NF-L levels were associated with longitudinal dementia status.

**DISCUSSION:** NF-L is a biomarker for neurodegeneration in DS, with potential for use in future clinical trials to prevent or delay dementia.

**Research in context:** *Systematic review:* The authors reviewed the literature using PubMed searches supplemented with our knowledge of pending papers in this research area. While blood NF-L has been associated with clinical features of progression in a number of neurodegenerative conditions, we have not identified any reports of NF-L associated with cognitive decline in DS, a genetic form of AD.

*Interpretation:* Our findings demonstrate the potential utility of NF-L as a blood biomarker of neurodegeneration in DS, a population that may not be able to tolerate more invasive procedures such as neuroimaging and lumbar punctures to track progression.

*Future directions:* The association between NF-L and other markers of longitudinal AD progression should be explored further in future work.

## 1. Introduction

Down syndrome (DS), caused by the trisomy, translocation, or partial trisomy of chromosome 21, is the most common genetic cause of intellectual disability (ID) with an estimated population of 6 million worldwide. Dementia is a common feature of the ageing process in DS due to the triplication of the amyloid precursor protein (APP) on chromosome 21 leading to brain pathology indicative of Alzheimer’s disease (AD) [1], a cumulative incidence for dementia in excess of 90% by the age of 65 [2], and a mean age at dementia diagnosis of 55 [3]. DS is therefore a genetic form of AD alongside autosomal dominant causes of AD [4]

Neurofilament light (NF-L) is one of the scaffolding cytoskeleton proteins of myelinated subcortical axons [5] and can now be reliably measured in blood using ultrasensitive single molecule array technology. Blood concentration of NF-L correlates well with corresponding CSF measures [6] and reflects axonal damage in neurological disorders, including frontotemporal dementia [7], multiple sclerosis [8], and familial and sporadic AD [9, 10]. NF-L correlates with other measures of disease stage and severity [10] but the utility of NF-L in populations with other genetic forms of AD is yet to be fully explored.

We aimed to explore the relationship between plasma NF-L levels, age, and dementia status in individuals with DS, as well as its independence from sex effects, premorbid intellectual ability levels, and longstanding epilepsy.

## 2. Methods

The North West Wales National Health Service (NHS) Research Ethics Committee provided ethical approval for a longitudinal study of cognitive ability and dementia in DS (13/WA/0194). For those with decision-making capacity written consent was obtained, while for those who did not have decision-making capacity a consultee indicated their agreement to participation.

Participants aged 16 and older were recruited across England via care homes, support groups, and local NHS sites. Participants with an acute physical or mental health condition were excluded until they had recovered; other details of the cohort have been previously described [11]. DS status was confirmed using DNA from saliva or blood and genotyped using Illumina OmniExpressExome arrays (San Diego, CA, USA); trisomy status was visually confirmed in GenomeStudio (see Table 1). APOE was determined using Thermo Fisher Scientific Taqman assays for rs7412 and rs429358 (Waltham, MA, USA).

**Table 1.** Participant demographics of all participants included in group and subgroup analyses

|  | **All participants** | **Dementia**<br>**(baseline)** | **No dementia**<br>**(baseline)** | **Participants with**<br>**follow up data** |
| --- | --- | --- | --- | --- |
| Number | 94 | 18 | 76 | 29 |
| Age at baseline<br>(mean ±<br>standard<br>deviation<br>(range)) | 42.68±14.87 (17-73) | 55.17±9.92 (40-69) | 39.72±14.34 (17-73) | 52.63±8.88 (40-72) |
| DS type | 89 (94.7%)<br>trisomy, 2 (2.1%)<br>translocation, 3 (3.2%) unknown | 18 (100.0%)<br>trisomy | 71 (93.4%)<br>trisomy, 2 (2.6%)<br>translocation, 3 (3.9%) unknown | 28 (96.6%)<br>trisomy, 1 (3.4%)<br>unknown |
| Sex | 41 (43.6%)<br>female, 53 (56.4%) male | 6 (33.3%)<br>female, 12 (66.7%) male | 35 (46.1%)<br>female, 41 (55.9%) male | 10 (34.5%)<br>female, 19 (65.5%) male |
| Ethnicity | 85 (90.4%)<br>white, 9 (9.6%)<br>other | 17 (94.4.0%)<br>white, 1 (5.6%)<br>other | 68 (89.5%)<br>white, 8 (10.5%)<br>other | 27 (93.1%)<br>white, 2 (6.8%)<br>other |
| Pre-dementia ID<br>level | 37 (39.4%) mild, 47 (50%)<br>moderate, 9 (9.6%) severe, 1 (1.2%) unknown | 6 (33.3%) mild, 9 (50.0%)<br>moderate, 3 (16.7%) severe | 31 (40.8%) mild, 38 (50%)<br>moderate, 6 (7.9%) severe, 1 (1.3%) unknown | 13 (44.8%) mild, 12 (41.4%)<br>moderate, 3 (10.3%) severe, 1 (3.4%) unknown |

|  |  |  |  |  |
| --- | --- | --- | --- | --- |
| APOE status | 68 (72.3%) non-APOE4 carrier, 23 (24.5%) APOE4 carrier, 3 (3.2%) unknown | 12 (66.7%) non-APOE4 carrier, 5 (27.8%) APOE4 carrier, 1 (5.5%) unknown | 56 (73.7%) non-APOE4 carrier, 18 (23.7%) APOE4 carrier, 2 (2.6%) unknown | 22 (75.9%) non-APOE4 carriers, 6 (20.7%) APOE4 carrier, 1 (3.4%) unknown |
| NFL level (median (range)) ng/L | 22.74 (6.11-136.91) | 63.76 (15.21-136.91) | 19.96 (6.11-116.84) | 32.67 (12.23 – 481.97) |

Assessment included a detailed interview with carers using the Cambridge Examination of Mental Disorders of Older People with Down’s syndrome (CAMDEX-DS) [12] to determine decline in several domains including memory. Premorbid ID level was defined according to the ICD10 diagnostic system’s descriptions of mild, moderate, and severe ID, based on carers’ reports of the individuals’ best ever level of functioning [13]. At baseline, dementia was defined as a confirmed clinical diagnosis. At follow-up, participants were classified according to whether they had retained or been given a diagnosis of dementia, or were being investigated for dementia.

Blood samples from 100 individuals were collected in lithium heparin tubes (Fisher, Loughborough, UK) and sent overnight for processing. Blood was layered over a similar amount of ficoll (GE Healthcare, Little Chalfont, UK), then centrifuged in a swing-out rotor for 40 minutes at 400g without brake. Plasma samples were stored at −80°C. Plasma NF-L concentration was measured by the same laboratory technician with reagents from a single lot using the Simoa NF-light^®^ assay (a digital sandwich immunoassay employing antibodies directed against the rod domain of NF-L) on an HD-1 Simoa analyzer, according to the protocol issued by the manufacturer (Quanterix, Lexington, MA, USA). Samples were run in duplicate and coefficients of variance (CV) for duplicates were set to be below 12%. All samples measured within the range spanned by the limits of quantification and inter-assay CV for the high and low concentration quality controls were 6.6 and 8.1% respectively.

All statistical tests were 2-sided and statistical significance was set at p<0.05. We tested associations between plasma NF-L samples and demographic or clinical factors using Mann-Whiney U, Kruskal-Wallis, and Spearman rank correlation tests as appropriate. Associations between plasma NF-L and dementia diagnosis were tested using logistic regression and log-transformed NF-L values, with adjustment for age and sex; we also adjusted for APOE4 status cross-sectionally.

## 3. Results

NF-L levels were obtained from 100 participants (age range 17-73); 5 results were excluded after failing to meet CV thresholds meaning 95 adults were included in subsequent analyses. Of adults aged 36 and older who are being targeted for longitudinal follow-up, 29/63 (46%) had completed a follow-up assessment at the time of this report (mean number of months between assessments 23.4, SD 3.9). One individual had suffered an occlusive cerebrovascular event 4-6 months prior to donating the blood sample and converted to dementia status at follow-up but was an outlier with an NF-L level of 481.97 ng/L, thus was excluded from cross-sectional analysis. For the remaining 94 individuals NF-L concentration had a median value of 22.74 ng/L, range 6.11-136.91 ng/L. At baseline, 18 of 94 participants had a clinical diagnosis of dementia (Table 1).

NF-L levels did not differ by premorbid ID level (Kruskal-Wallis test, p=0.195), sex (Mann-Whitney U test, p=0.837) or longstanding epilepsy (Mann-Whitney U test, p=0.858). NF-L level and age of participants were significantly correlated (Spearman’s rho = 0.789, p<0.001) (Figure 1), such that those aged 35 and older had significantly higher levels of NF-L compared to younger individuals (median 11.52 vs. 32.42, Mann-Whitney U test p<0.001). Those with dementia had significantly higher levels of NF-L (median 63.76 ng/L vs. 19.96 ng/L; Mann-Whitney U test p<0.001), and a logistic regression model adjusting for age, sex, and APOE4 status revealed that NF-L levels remained predictive of dementia status (p=0.022).

**Figure 1.**
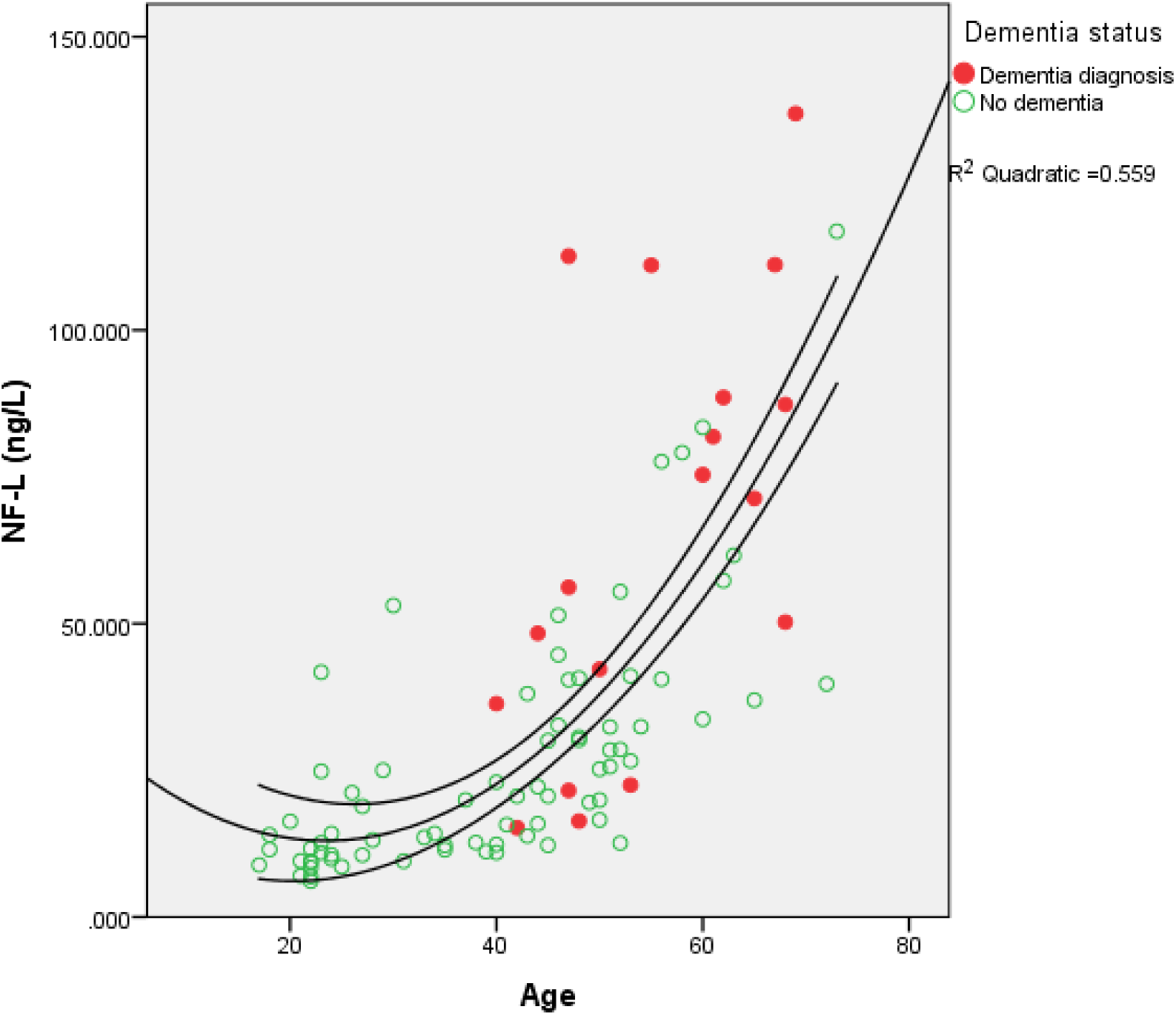
NF-L concentration by age of individuals with DS.

Seven (24.1%) of 29 individuals with follow-up cognitive data had a clinical diagnosis of dementia at baseline, all with the dementia diagnosis retained at follow-up, with a further two (6.9%) individuals converting to dementia status by follow-up, while three (10.3%) participants were under investigation for dementia at follow-up. Predictive validity of NF-L levels at baseline was explored by combining individuals with confirmed or suspected dementia at follow-up (n=12, median NF-L 77.38 ng/L), and comparing them against those who remained dementia-free (median NF-L 19.94 ng/L). Higher levels of NF-L at baseline predicted the likelihood of dementia at follow-up, even when adjusted for age and sex (p=0.036).

## 4. Discussion

We demonstrated that NF-L measured in blood using an ultra-sensitive assay is strongly associated with age and dementia status in individuals with DS, and baseline levels were predictive of dementia diagnosis over time. Furthermore, NF-L levels did not differ according to severity of premorbid ID or by longstanding epilepsy diagnosis (a common neurological comorbidity in DS), suggesting that it is a stable and feasible biomarker that can be used in clinical populations.

Our results indicate that this marker could pinpoint onset of neurodegeneration in DS. NF-L showed an age relationship in keeping with post-mortem data and amyloid positron emission tomography (PET) studies of AD pathology in adults with DS [1, 14]. Although NF-L is a marker of axonal damage, and thus not specific to AD [7-9, 15], in a genetically predisposed population such as DS where AD is almost always the cause of dementia, the lack of specificity is arguably less of an issue and NF-L could potentially be used as a biomarker for treatment response. Normalization of serum/plasma NF-L in response to treatment has already been demonstrated in patients with multiple sclerosis [8, 16].

Although further work is required to establish long-term predictive and concurrent validity of NF-L, our data suggests that this biomarker could be instrumental in allowing an experimental medicine approach in DS and other high-risk populations to test treatments that might prevent or delay dementia onset.

## Acknowledgements

The authors would like to thank all the participants in this study for their time.

This work was funded by a Wellcome Trust Strategic Award (grant number: 098330/Z/12/Z) conferred upon The London Down Syndrome (LonDownS) Consortium. The funder had no role in study design; the collection, analysis, or interpretation of the data; the writing of the report; or the decision to submit the report for publication.

HZ is a Wallenberg Academy Fellow and receives support from the European Research Council, the Swedish Research Council, and Frimurarestiftelsen. HZ is further supported by the Wolfson Foundation, the Wellcome Trust, and the UK Dementia Research Institute. DN is also supported by the National Medical Research Council Singapore (NMRC/CIRG/1438/2015) and Singapore Ministry of Education AcRF-Tier2 (2015-T2-1-023). This research was further supported by the National Institute for Health Research networks (mental health, dementias and neurology) and participating NHS trusts. We would like to thank our NHS network of sites that helped to identify participants.

The LonDownS Consortium principal investigators are Andre Strydom (chief investigator), Department of Forensic and Neurodevelopmental Sciences, Institute of Psychiatry, Psychology and Neuroscience, King’s College London, London, UK, and Division of Psychiatry, University College London, London, UK; Elizabeth Fisher, Department of Neurodegenerative Disease, UCL Institute of Neurology, London, UK; Dean Nizetic, Blizard Institute, Barts and the London School of Medicine, Queen Mary University of London, London, UK, and Lee Kong Chian School of Medicine, Nanyang Technological University, Singapore, Singapore; John Hardy, Reta Lila Weston Institute, Institute of Neurology, University College London, London, UK, and UK Dementia Research Institute at UCL, London, UK; Victor Tybulewicz, Francis Crick Institute, London, UK, and Department of Medicine, Imperial College, London, UK; and Annette Karmiloff-Smith (Birkbeck University of London, London, UK, deceased). Other members of The LonDownS Consortium who contributed to data collection are Sarah Hamburg and Rosalyn Hithersay (both Department of Forensic and Neurodevelopmental Sciences, Institute of Psychiatry, Psychology and Neuroscience, Kings College London, London, UK, and Division of Psychiatry, University College London, London, UK).

Author contributions
Conception and design of the study: AS, HZ; acquisition of data: AH, KM, JH, JG, DN, HZ; analysis of data: AS, CS; writing the manuscript: AS, CS; revising the manuscript for important intellectual content: all co-authors.

